# The effect of carbamazepine on bone structure and strength in control and Osteogenesis Imperfecta (*Col1a2* ^+/*p.G610C*^) mice

**DOI:** 10.1101/2022.02.23.481692

**Authors:** Martha Blank, Narelle E McGregor, Lynn Rowley, Louise HW Kung, Blessing Crimeen-Irwin, Ingrid J Poulton, Emma C Walker, Jonathan H Gooi, Shireen R. Lamande, Natalie A Sims, John F. Bateman

## Abstract

The inherited brittle bone disease osteogenesis imperfecta (OI) is commonly caused by *COL1A1* and *COL1A2* mutations that disrupt the collagen I triple helix. This causes intracellular endoplasmic reticulum (ER) retention of the misfolded collagen and can result in a pathological ER stress response. A therapeutic approach to reduce this toxic mutant load could be to stimulate mutant collagen degradation by manipulating autophagy and/or ER-associated degradation. Since carbamazepine (CBZ) both stimulates autophagy of misfolded collagen X and improves skeletal pathology in a metaphyseal chondrodysplasia model, we tested the effect of CBZ on bone structure and strength in 3 week-old male OI *Col1a2* ^*+/p*.*G610C*^ and control mice. Treatment for 3 or 6 weeks with CBZ, at the dose effective in metaphyseal chondrodysplasia, provided no therapeutic benefit to *Col1a2* ^*+/p*.*G610C*^ mouse bone structure, strength, or composition, measured by micro-computed tomography, three point bending tests and Fourier-transform infrared microspectroscopy. In control mice however, CBZ treatment for 6 weeks impaired femur growth and led to lower femoral cortical and trabecular bone mass. These data, showing the negative impact of CBZ treatment on the developing mouse bones, raise important issues which must be considered in any human clinical applications of CBZ in growing individuals.

## Introduction

The inherited brittle bone disease osteogenesis imperfecta (OI) is commonly caused by mutations that compromise the biosynthesis or structure of type I collagen, the predominant organic structural component of bone. The two collagen I α-chain subunits, α1(I) and α2(I), which form the [α1(I)]_2_α2(I) collagen protein trimers are encoded by *COL1A1* and *COL1A2*. Autosomal dominant mutations in these genes cause more than 85% of OI cases ^1,2^, and fall into two broad functional groups. Mutations that reduce collagen I expression, such as nonsense mutations which lead to mRNA decay, result in milder clinical phenotypes (e.g., OI type I), whereas mutations introducing missense changes causing structural disruptions in collagen proα-chains lead to more severe OI phenotypes (e.g. OI types II, III, IV). These severe phenotypes most commonly exhibit impaired collagen I triple helix structure and stability, usually caused by glycine substitutions that interrupt the obligatory Gly-X-Y amino acid repeat sequence required for collagen helix folding. A common consequence is dysregulated collagen proteostasis, with intracellular retention and aggregation of the misfolded collagen ^3^. Its intracellular accumulation can induce an endoplasmic reticulum (ER) stress response which might contribute to the pathology ^2,4^.

A mouse model of OI type IV (*Col1a2* ^*+/p*.*G610C*^) with a Gly to Cys *Col1a2* missense mutation in the triple helix, based on an Old Order Amish OI mutation ^5^, provides us with an experimental tool for pre-clinical therapeutic testing. Previous studies on this mouse model have shown that, as expected, the glycine substitution mutation leads to intracellular mutant collagen accumulation, a poorly characterized non-canonical ER stress response and downstream impairment in osteoblast differentiation and function ^6^ with clinically-relevant changes to bone structure and fragility ^7^.

While the molecular detail of how helix-disrupting OI mutations cause intracellular ER stress and bone pathology are not fully resolved ^8^, a promising therapeutic approach would be to reduce the load of toxic misfolded collagen and thereby reduce the OI bone pathology to a milder phenotype. There are few FDA-approved drugs that could be re-purposed to achieve this. Rapamycin, which stimulates autophagy via the mTOR pathway, showed promise on osteoblasts from the *Col1a2* ^*+/p*.*G610C*^ mouse *in vitro* ^6^. However, *in vivo* testing in the *Col1a2* ^*+/p*.*G610C*^ mouse demonstrated that while some OI bone trabecular properties were improved, off-target effects suppressing longitudinal and transverse bone growth in OI mice precluded rapamycin as a viable therapeutic option ^9^. Another drug stimulating mTOR-independent autophagy, carbamazepine (CBZ), has been effective in a mouse model of metaphyseal chondrodysplasia, type Schmid (MCDS), where intracellular accumulation of misfolded mutant collagen X causes ER stress and downstream pathological signalling ^10^. CBZ-stimulated degradation reduced the toxic mutant collagen load and ER stress, and improved the MCDS cartilage and bone pathology ^11,12^. As a result CBZ is now in clinical trials with MCDS patients (mcds-therapy.eu). Because of the broad similarity of the molecular phenotypes in OI and MCDS (collagen misfolding and intracellular retention) and the predicted therapeutic effect of CBZ in stimulating autophagic clearance of misfolded collagen I, there is considerable interest in CBZ as a potential OI therapy.

Here we treated 3 week old *Col1a2* ^*+/p*.*G610C*^ and control mice with CBZ for 3 and 6 weeks from weaning and assessed structure and strength of the long bones. While CBZ treatment for 3 weeks does not affect control bone, longer term treatment (6 weeks) during this period of active bone growth reduces bone length, width, and strength. Furthermore, CBZ treatment for 3 or 6 weeks does not rescue the OI bone phenotype in *Col1a2* ^*+/p*.*G610C*^ mice.

## Materials and Methods

Heterozygous α2(I)-G610C mice (*Col1a2* ^*+/p*.*G610C*^) ^5^ were obtained from Jackson Laboratory, Bar Harbor, ME, USA (B6.129(FVB)-Col1a2^tm1Mcbr/J^; stock 007248) and maintained on a C57BL/6J background to generate heterozygous *Col1a2* ^*+/p*.*G610C*^ and control experimental mice. Mice husbandry and health monitoring is previously described ^9^. All procedures were approved by the Murdoch Children’s Research Institute Animal Ethics Committee (#A798). Male control and *Col1a2* ^*+/p*.*G610C*^ mice were treated by oral gavage with CBZ (250 mg/kg/day) from 3 to 6 weeks ^11,12^. A second cohort was additionally treated from 6 to 9 weeks by subcutaneous implantation of a slow-release pellet of carbamazepine (250 mg/kg/day) (Innovative Research of America USA C-113) ^11^. At 6 and 9 weeks, mice were euthanased and tissues harvested.

### Micro-computed tomography (μCT)

Femora were assessed on a Skyscan 1276 micro-CT system (Bruker, Aartselaar, Belgium) at 9 micron voxel resolution as previously described ^9^, using NRecon (version 1.7.1.0), DataViewer (version 1.5.4), and CT Analyzer (version 1.16.4.1). Trabecular bone structure, and multi-level thresholding were carried out in the metaphysis, starting at 7.5% of bone length from the distal femoral growth plate and along 15% of total bone length. Cortical structure was assessed in the diaphysis, starting at 30% of bone length from the distal femoral growth plate and along 15% of total bone length. Medio-lateral widths and cranio-caudal widths were determined at the femoral midpoint. Single thresholds for cortical bone were 0.487 g/cm^3^ and 0.786 g/cm^3^ CaHA for 6 and 9 week old mice, respectively and for trabecular bone 0.219 g/cm^3^ and 0.435 g/cm^3^ CaHA for 6 and 9 week old mice, respectively. To segregate bone into multiple density levels, nonparametric unsupervised 4 level Otsu-thresholding algorithm was used, as previously described ^13^. Each density area was normalised to total bone area and thresholds of 0.301-0.613, 0.614-1.003 and >1.003 mg/cm^2^ CaHA were used for low-, mid-, and high-density bone areas, respectively; the background level (0-0.3 mg.cm^2^ CaHA) was discarded.

### Three point bending strength tests

After μCT, femora were stored in 70% ethanol. Three point bending was performed with a Bose Biodynamic 5500 Test instrument (Bose, DE, USA) and WinTest 7 software. Femora were centred onto supporting pins with a span width of 6 mm and loaded with 0.5 mm/s. As described previously ^14^, the load/deformation data was collected over 5 seconds with a sampling rate of 250 Hz. For material properties, the data was normalized to medio-lateral and cranio-caudal widths and average cortical thickness of the diaphysis of each bone.

### Synchrotron-based Fourier transform infrared microspectroscopy (sFTIRM)

sFTIRM was conducted on a Bruker Hyperion 2000 IR microscope coupled to a V80v FTIR spectrometer at the Australian Synchrotron IR Microspectroscopy beamline, as previously described ^15^. Briefly, 2 μm undecalcified methyl methacrylate (MMA) embedded tibial sections ^16^ were placed on 0.5 mm barium fluoride windows (Crystan Limited, UK), and 16×16 μm regions of interest were measured with 10 μm spacing from the periosteal edge at 1.5 mm proximal to the growth plate in the medial cortex. Spectra were collected using a wideband detector with 256 co-added scans per pixel spectral resolution in transmission mode. For each sample background spectra were collected through both clear barium fluoride and MMA. Data acquisition and analysis was performed with Bruker OPUS version 8.0. To analyse the raw spectra, each spectrum was corrected for water vapor, baseline corrected at 1800,1200 and 800 cm^-1^ and absorbed MMA was subtracted. Bone composition was determined by integrating areas as follows: mineral:matrix ratio using amide I (1180-916:1712-1588cm^-1^) or amide II (1180-916:1600-1500cm^-1^), carbonate substitution (890-852:1180-916 cm^-1^) and collagen compaction (1712-1588:1600-1500cm^-1^), and crystallinity from 1030-1020cm^-1^, all as previously described ^15,17^.

### Statistical analysis

All graphs show mean ± SEM. Sample number (n) is indicated in figure legends. Statistical significance was calculated with GraphPad Prism 9 (version 9.1.2.) by two-way ANOVA and Tukey’s multiple comparison test for all comparisons except for the regional analysis for sFTIRM and Otsu thresholding where two-stage linear step-up procedure of Benjamini, Krieger and Yekutieli was used.

## Results

Six week old *Col1a2* ^*+/p*.*G610C*^ mice have defective bone structure that is not modified by CBZ

### treatment from 3 weeks of age

Previous studies have shown that from 8 weeks of age (but not at 10 days of age), *Col1a2* ^*+/p*.*G610C*^ mice have less cortical and trabecular bone and lower bone strength than controls ^5,7,9,18^. We showed that the phenotype can also be detected at 6 weeks of age. Femora from 6 week old vehicle-treated *Col1a2* ^*+/p*.*G610C*^ mice were narrower with significantly lower cortical area, periosteal and endocortical perimeters compared to controls (Figure 1A-D). In addition, the marrow area, mean polar moment of inertia and medio-lateral and cranio-caudal widths were significantly lower than controls (Table S1). *Col1a2* ^*+/p*.*G610C*^ femora from 6 week old mice also had significantly lower trabecular bone volume, number, separation, and shorter femora than controls (Figure 1E-I). Treatment with 250mg/kg/day CBZ for 3 weeks, a dose equivalent to that used in treating epilepsy in human subjects taking into account species differences ^19^ and to correct cartilage defects in the MCDS mouse ^11,12^, did not significantly modify cortical or trabecular bone mass in either control or *Col1a2* ^*+/p*.*G610C*^ femora (Figure 1A-I and Table S1). CBZ from 3 to 9 weeks of age had no effect in

**Figure 1.**
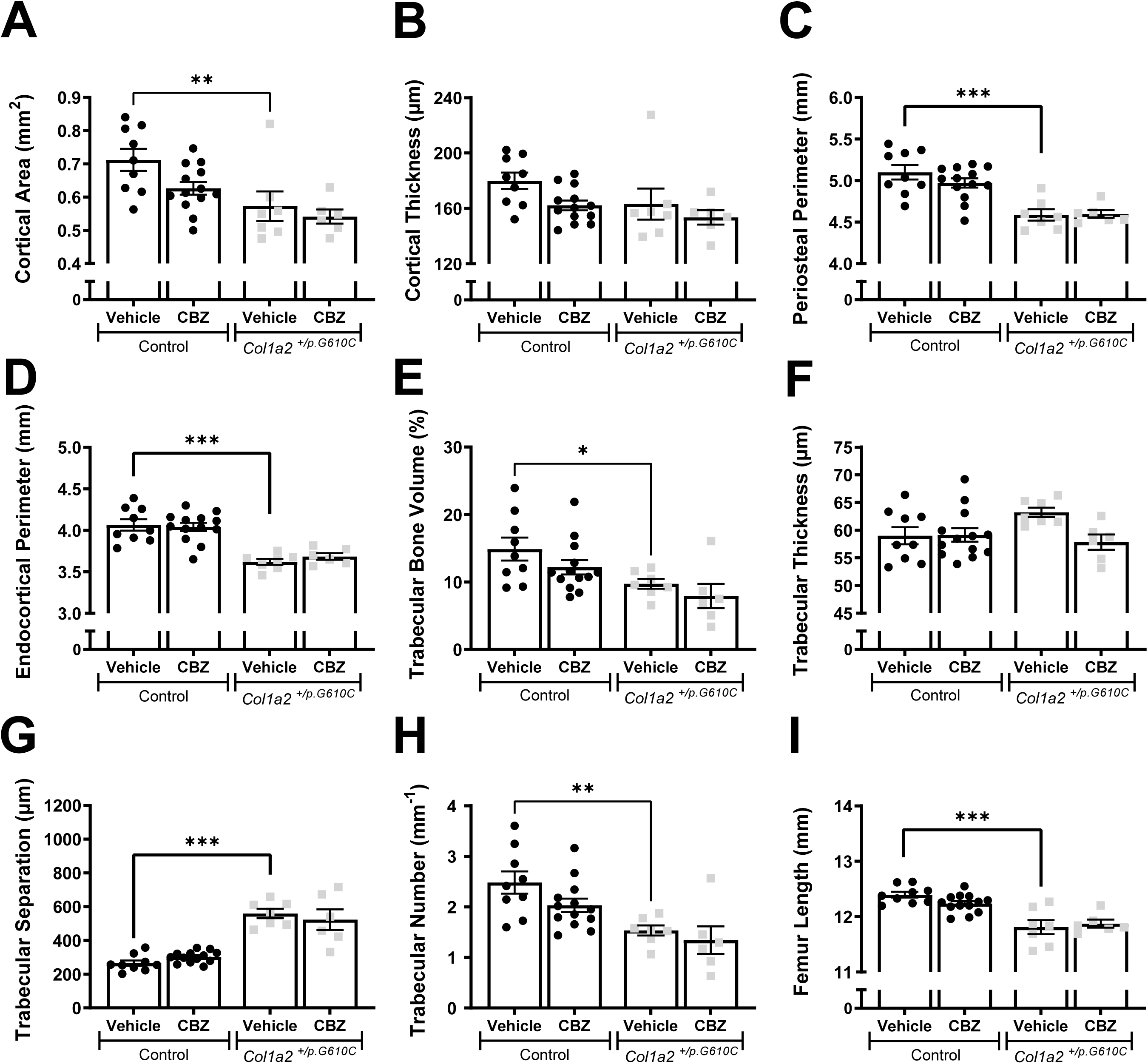
Femoral bone structure of 6 week old male control and *Col1a2* ^*+/p*.*G610C*^ mice treated with vehicle or carbamazepine (CBZ) for 3 weeks assessed by micro-computed tomography (μCT). Shown are (A) cortical area, (B) cortical thickness, (C) periosteal perimeter and (D) endocortical perimeter, (E) trabecular bone volume, (F) trabecular thickness, (G) trabecular separation, (H) trabecular number and (I) femur length from male control and *Col1a2* ^*+/p*.*G610C*^ mice. Data shown are mean ± SEM, n= 6-13 mice/group. * p<0.05, ** p<0.01, *** p<0.001 vs. treatment-matched controls.

### *Col1a2* ^*+/p*.*G610C*^ mice but impaired bone growth in control mice

The lower trabecular bone mass, and smaller femoral length and width of 6 week old vehicle-treated *Col1a2* ^*+/p*.*G610C*^ mice compared to controls were also detected at 9 weeks of age (Figure 2 and Table S2). The extended period of CBZ treatment (from 3 to 9 weeks of age) had no significant effect on trabecular or cortical structure in *Col1a2* ^*+/p*.*G610C*^ femora (Figure 2 and Table S2). Although short-term CBZ treatment in control mice for 3 weeks had no effect on bone structure (Figure 1 and Table S1), a longer duration of CBZ treatment for 6 weeks led to significantly lower cortical area, cortical thickness, periosteal and endocortical perimeters (Figure 2A-D), marrow area, mean polar moment of inertia, medio-lateral and cranio-caudal widths (Table S2), indicating suppressed transverse bone growth. Six weeks of CBZ treatment in control mice lowered trabecular bone volume, thickness, and number (Figure 2E, F, H). In these 9 week old mice, trabecular thickness and number were lower than in the younger vehicle-treated control mice (Figure 1E, F, H), suggesting loss of trabecular bone mass, rather than impaired trabecular bone accrual. In addition, while femoral length was greater at 9 weeks than 6 weeks in vehicle-treated control mice, CBZ-treated control mice had shorter femora than vehicle-treated controls (Figures 1I and 2I), indicating that a longer duration of CBZ treatment suppressed both transverse and longitudinal bone growth.

**Figure 2.**
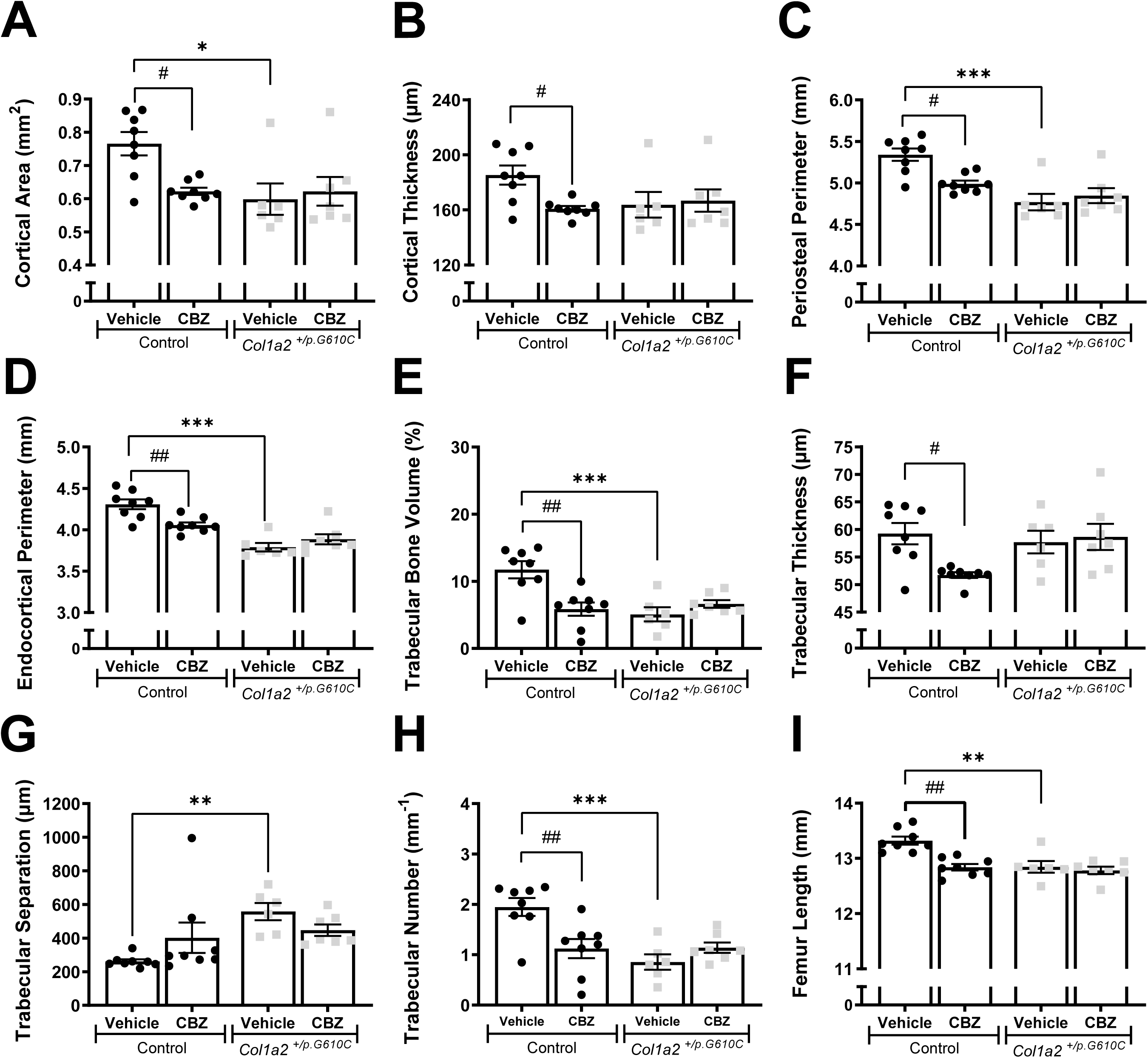
Femoral bone structure of 9 week old male control and *Col1a2* ^*+/p*.*G610C*^ mice treated with vehicle or carbamazepine (CBZ) for 6 weeks assessed by micro-computed tomography (μCT). Shown are (A) cortical area, (B) cortical thickness, (C) periosteal perimeter and (D) endocortical perimeter, (E) trabecular bone volume, (F) trabecular thickness, (G) trabecular separation, (H) trabecular number and (I) femur length from male control and *Col1a2* ^*+/p*.*G610C*^ mice. Data shown are mean ± SEM, n= 6-8 mice/group. * p<0.05, ** p<0.01, *** p<0.001 vs. treatment-matched controls. # p<0.5, ## p<0.01 vs. genotype-matched controls.

### *Col1a2* ^*+/p*.*G610C*^ mice have disrupted cortical bone consolidation, but this is modified by CBZ only in control mice

Cortical bone consolidates by changing from a material with a high proportion of low-density bone to a material with mainly high-density bone ^16^. This transition can be measured along the length of the growing bone from the growth plate (more low density bone) to the diaphysis (more high density bone). Since cortical diaphyseal bone of *Col1a2* ^*+/p*.*G610C*^ mice has greater average tissue mineral density ^18^, we used multi-level thresholding to assess whether high density material accrued more rapidly in less mature regions of the femoral metaphysis of *Col1a2* ^*+/p*.*G610C*^ mice.

When total proportions of low-, mid-and high-density bone were assessed in vehicle-treated animals, *Col1a2* ^*+/p*.*G610C*^ femora had a higher proportion of mid-density bone material than controls, with no significant difference in proportions of low-or high-density bone (Figure 3). This was not modified by CBZ treatment (Figure 3).

**Figure 3.**
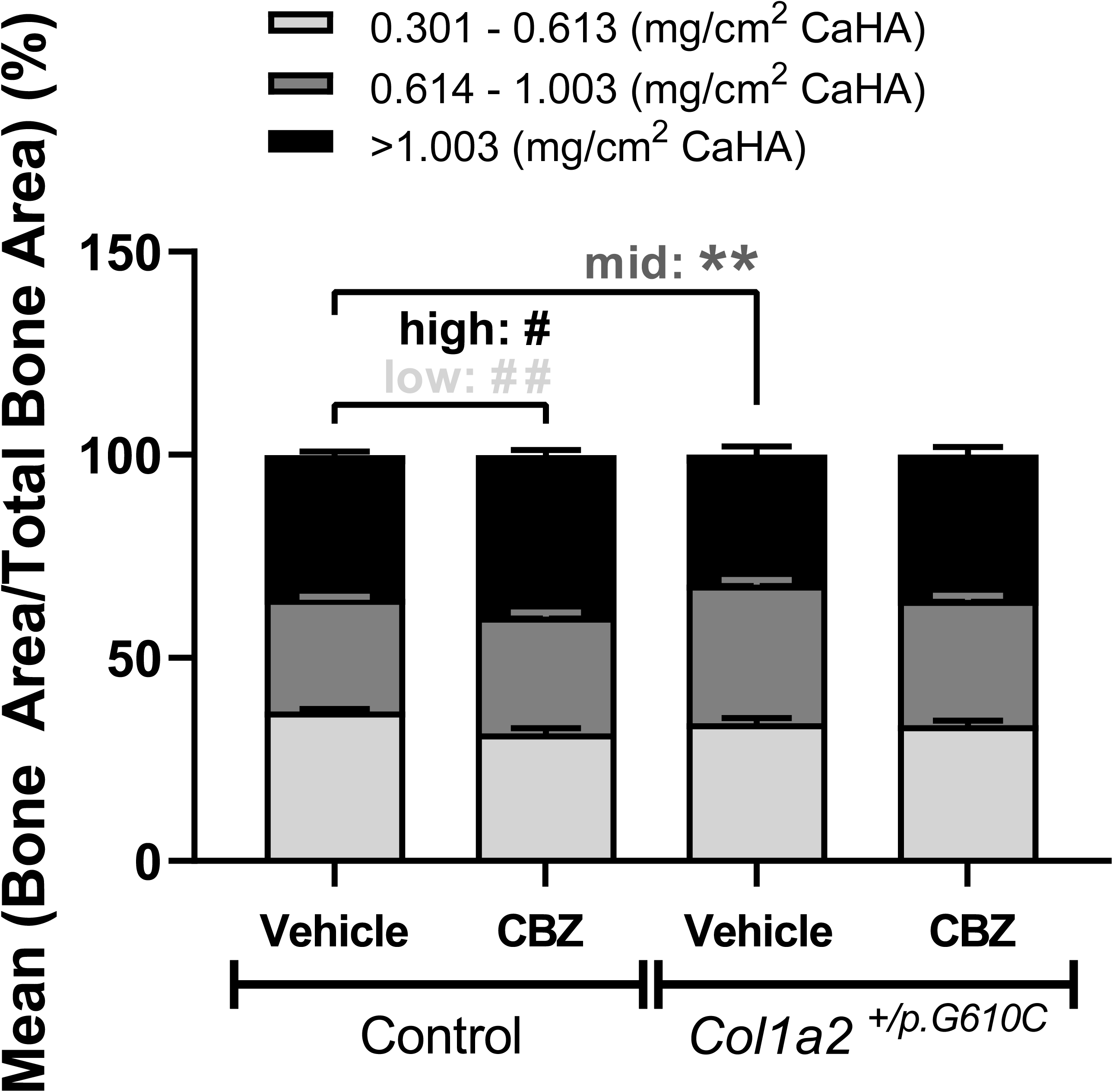
Multi-level-thresholding analysis of the average femoral metaphysis segregated by low-(light-grey), mid-(dark-grey) and high-density (black) levels of the femoral cortex from 9 week old male control and *Col1a2* ^*+/p*.*G610C*^ mice treated with vehicle or carbamazepine (CBZ) for six weeks by micro-computed tomography (μCT). Data are mean ± SEM; n= 5-8 mice/group. ** p<0.01 vs. treatment-matched controls. # q<0.05, ## q<0.01 vs. treatment- and region-matched controls.

When assessed along the length of the bone, vehicle-treated control and *Col1a2* ^*+/p*.*G610C*^ bones both showed the expected reduction in low-density and increase in high-density bone with increasing distance from the growth plate (Supplementary Figure 1A). However, *Col1a2* ^*+/p*.*G610C*^ bones had less high density bone in the least mature region (close to the growth plate), more low density bone in the most mature region (near the diaphysis), and a higher proportion of mid-density bone along the full length of the metaphysis, indicating region-specific differences in cortical bone maturation in this model of OI.

CBZ treatment from 3 to 9 weeks of age increased the proportion of high-density bone and reduced the proportion of low-density bone in control mice (Figure 3). This was significant along the full extent of the metaphysis, indicating that it existed at all stages of cortical bone consolidation and that CBZ could change the type of bone deposited (Supplementary Figure 1B). CBZ treatment did not affect these parameters in *Col1a2* ^*+/p*.*G610C*^ femora at any point along the metaphysis (Supplementary Figure 1C).

### sFTIRM shows mineral:matrix content of *Col1a2* ^*+/p*.*G610C*^ bone progressively increases in deeper sub-periosteal bone

Eight week old, but not 10 day old, male *Col1a2* ^*+/p*.*G610C*^ mice exhibit high mineral:matrix ratio ^7^, consistent with an imbalance between collagen production and mineral accrual ^20^. Since 12 week old tibial cortex exhibits a gradient of matrix maturation from the periosteal edge moving inwards, including mineral accrual, collagen compaction and carbonate substitution ^15,17^, we assessed whether the OI phenotype or CBZ treatment modified bone matrix composition and maturation.

Both 6 and 9 week old vehicle-treated *Col1a2* ^*+/p*.*G610C*^ tibial cortex had greater mineral:matrix ratio (higher phosphate:amide I ratio) than controls (Figure 4A, C), indicating that this matrix defect is already significant at 6 weeks of age. In vehicle-treated animals, the mineral:matrix ratio increased in both control and *Col1a2* ^*+/p*.*G610C*^ bone at 10 μm and 20 μm inwards from the periosteal edge compared to the periosteum, at both 6 and 9 weeks of age; the slope of this increase was notably greater in the more mature 9 week old bone (Figure 4B, D). Additionally, 6 week old *Col1a2* ^*+/p*.*G610C*^ bones had a higher mineral:matrix ratio at 10 μm and, in 9 week old mice, at both 10 μm and 20 μm from the periosteum compared to the same regions in control bones (Figure 4B, D). The mineral:matrix ratio using the amide II reference showed a similar response (Table 1 and Figure S2A, E). This indicates that although mineral is initially deposited at normal levels on the growing periosteum of *Col1a2* ^*+/p*.*G610C*^ bones, it accrues mineral more rapidly in *Col1a2* ^*+/p*.*G610C*^ mice than in controls.

**Figure 4.**
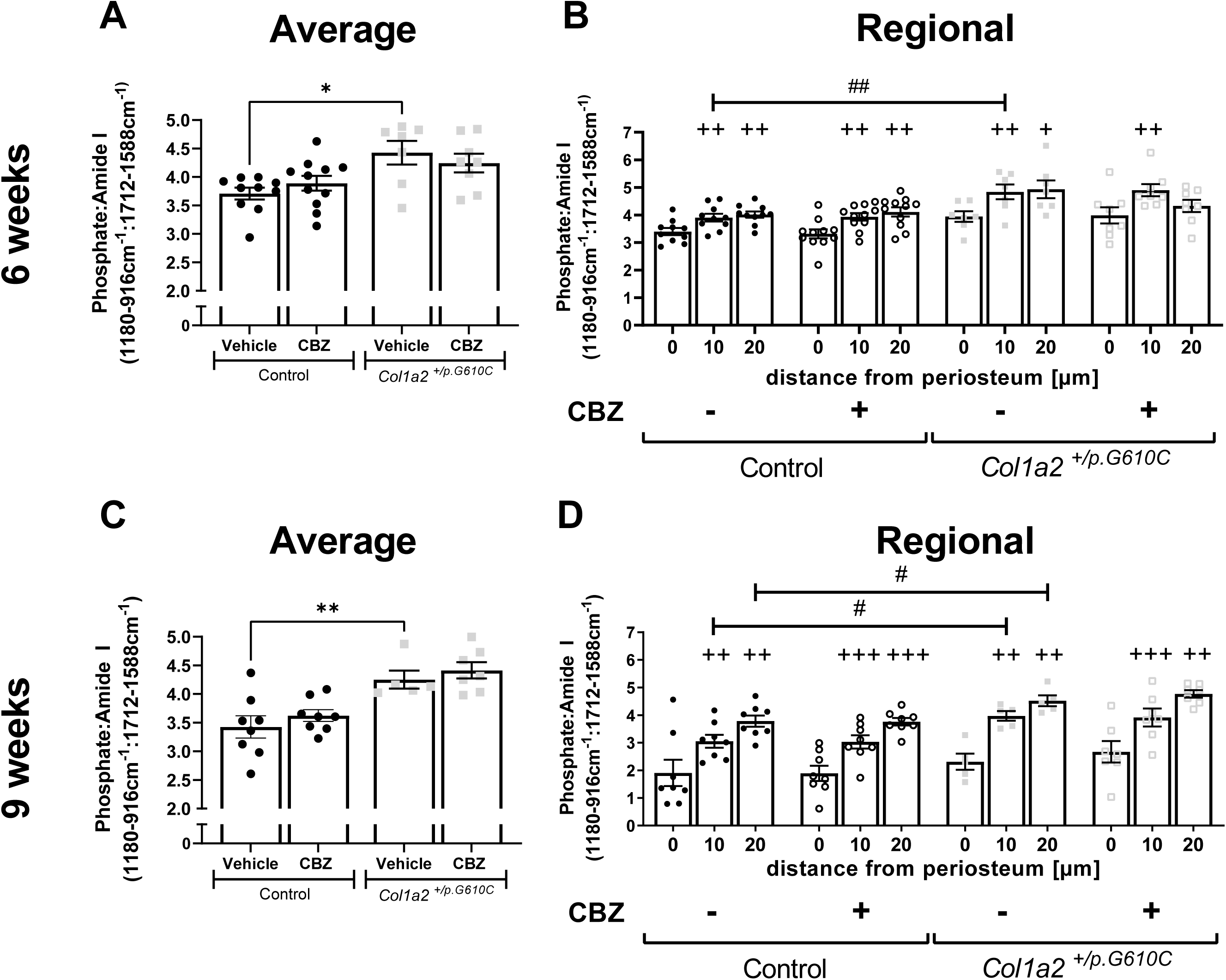
Analysis of average (A,C) and regional (B,D) mineral:matrix ratios of tibial cortex from 6 week (A,B) and 9 week (C,D) old male control and *Col1a2* ^*+/p*.*G610C*^ mice treated with vehicle or carbamazepine (CBZ) for three or six weeks, respectively by synchrotron Fourier transform infrared microspectroscopy (sFTIRM). Data is shown as the average phosphate:amide I ratio and regional measurements, which show data at 10 micron intervals with increasing distance from the periosteum. Ratios were calculated from integrated areas of phosphate (1180-916cm^-1^), amide I (1712-1588cm^-1^) curves. Data are mean ± SEM; n=7-11 mice/group for 6 week old mice and n= 5-8 mice/group for 9 week old mice. * p<0.05, ** p<0.01 vs. treatment-matched controls. + q<0.05, ++ q<0.01, +++ q<0.001 vs. genotype- and treatment-matched region 0 μm. # q<0.05, ## q<0.01 vs. treatment- and region-matched controls.

**Table 1.** Additional bone composition parameters of tibial cortex from 6 week and 9 week old male control and *Col1a2* ^*+/p*.*G610C*^ mice treated with vehicle or carbamazepine (CBZ) for three and six weeks respectively by synchrotron Fourier transform infrared microspectroscopy (sFTIRM). Ratios were calculated from integrated areas of phosphate (1180-916cm^-1^), carbonate (890-852cm^-1^), amide I (1712-1588cm^-1^) and amide II (1600-1500cm^-1^) curves. Crystallinity sub-peak was calculated by the integrated area from 1030-1020cm^-1^. Data shown are mean ± SEM; n= 7-11 mice/group for 6 week old mice and n= 5-8 mice/group for 9 week old mice. * p<0.05, ** p<0.01, *** p<0.01 vs. controls.

| **6 weeks** |  |  |  |  |
| --- | --- | --- | --- | --- |
|  | **Control** |  | ***Col1a2*<sup>+/-p.G610C</sup>** |  |
|  | Vehicle<br>(n= 10) | CBZ<br>(n= 11) | Vehicle<br>(n= 7) | CBZ<br>(n= 8) |
| Phosphate:Amide II | 6.49 ± 0.13 | 6.49 ± 0.08 | 7.52 ± 0.39 ** | 7.22 ± 0.20 |
| Carbonate:Phosphate | 0.009 ± 0.0003 | 0.008 ± 0.0002 | 0.007 ± 0.0004 * | 0.008 ± 0.0002 |
| Crystallinity | 0.72 ± 0.03 | 0.72 ± 0.04 | 0.86 ± 0.06 | 0.76 ± 0.05 |
| Amide I:II | 1.78 ± 0.02 | 1.72 ± 0.04 | 1.72 ± 0.04 | 1.74 ± 0.03 |
| **9 weeks** |  |  |  |  |
|  | **Control** |  | ***Col1a2*<sup>+/-p.G610C</sup>** |  |
|  | Vehicle<br>(n= 8) | CBZ<br>(n= 8) | Vehicle<br>(n= 5) | CBZ<br>(n= 7) |
| Phosphate:Amide II | 5.61 ± 0.21 | 6.12 ± 0.17 | 7.14 ± 0.30 *** | 7.01 ± 0.16 |
| Carbonate:Phosphate | 0.009 ± 0.0005 | 0.009 ± 0.0001 | 0.009 ± 0.0002 | 0.009 ± 0.0004 |
| Crystallinity | 0.17 ± 0.03 | 0.14 ± 0.03 | 0.13 ± 0.03 | 0.12 ± 0.02 |
| Amide I:II | 1.72 ± 0.04 | 1.76 ± 0.02 | 1.73 ± 0.03 | 1.65 ± 0.03 |

Average crystallinity and amide I:II ratio (collagen compaction) were not significantly different between vehicle-treated control and *Col1a2* ^*+/p*.*G610C*^ bone at 6 or 9 weeks of age (Table 1). Although carbonate:phosphate ratio was lower in vehicle-treated *Col1a2* ^*+/p*.*G610C*^ bones than control at 6 weeks of age (Table 1), this genotype-dependent effect was not observed at 9 weeks of age. While average carbonate:phosphate ratio was lower in vehicle-treated *Col1a2* ^*+/p*.*G610C*^ than controls at 6 weeks of age, region-specific analysis showed no change in carbonate:phosphate ratios with matrix maturation in either controls or *Col1a2* ^*+/p*.*G610C*^ bones at any age (Figure S2B, F). However, average crystallinity increased and amide I: II ratio decreased with increasing distance from the periosteum in both genotypes at 9 weeks of age (Figure S2G, H), suggesting that *Col1a2* ^*+/p*.*G610C*^ bone undergoes normal mineral maturation at 9 weeks of age, apart from the abnormal increase in mineral relative to collagen. No significant effects of CBZ were detected by sFTIRM in either control or *Col1a2* ^*+/p*.*G610C*^ mice (Figure 4, Figure S2, Table 1).

### CBZ does not improve the *Col1a2* ^*+/p*.*G610C*^ bone strength defect but reduces bone strength in control mice

Three point bending tests in vehicle-treated mice showed that femora from 6 and 9 week old *Col1a2* ^*+/p*.*G610C*^ mice were weaker than control femora (Figure 5). This difference was greater in 9 week old mice, which showed a ∼60 % lower maximum load than controls (Figure 5E) while 6 week old *Col1a2* ^*+/p*.*G610C*^ femora were only ∼30 % weaker than their age-matched controls (Figure 5A). *Col1a2* ^*+/p*.*G610C*^ femora were also more brittle compared to control femora, and again this was more pronounced in 9 week old mice, shown by a ∼90 % lower post yield displacement (Figure 5F). There were also greater differences between the genotypes in yield load (Table S1, S2) in 9 week old than 6 week old mice.

**Fig 5.**
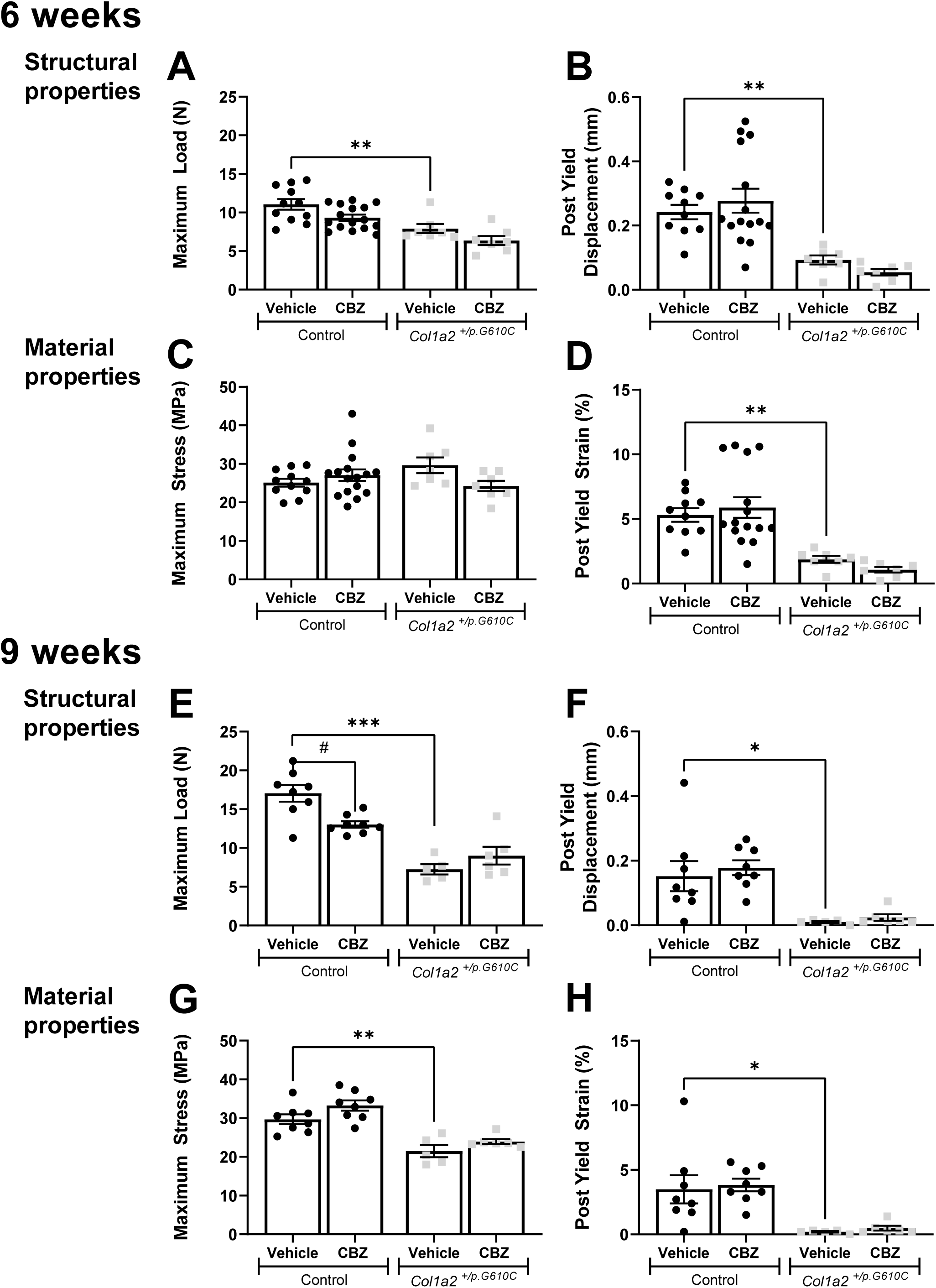
Results of three point bending tests, including raw data (structural properties) and those corrected for bone size (material properties) of femora from 6 week and 9 week old male control and *Col1a2* ^*+/p*.*G610C*^ mice treated with vehicle or carbamazepine (CBZ) for three and six weeks respectively. Shown are (A) maximum load, (B) post yield displacement, (C) maximum stress and (D) post yield strain of femora from 6 week old control and *Col1a2* ^*+/p*.*G610C*^ mice and (E) maximum load, (F) post yield displacement, (G) maximum stress and (H) post yield strain of femora from 9 week old control and *Col1a2* ^*+/p*.*G610C*^ mice. Data shown are mean ± SEM, n= 7-16 mice/group for 6 week old mice and n= 6-8 mice/group for 9 week old mice. * p<0.05, ** p<0.01, *** p<0.001 vs. treatment-matched controls. # p<0.05 vs. genotype-matched controls.

Since *Col1a2* ^*+/p*.*G610C*^ femurs were narrower than controls, strength testing data was corrected for bone widths and cortical thickness. When corrected for bone size, vehicle-treated *Col1a2* ^*+/p*.*G610C*^ femora were still more brittle than controls at both ages, shown by a lower post yield strain (Figure 5D, H), and lower elastic deformation, shown by a lower yield strain (Table S1, S2). The stiffness of *Col1a2* ^*+/p*.*G610C*^ femora was double that of controls, shown by a higher elastic modulus (Table S2). There was no difference in maximum stress in *Col1a2* ^*+/p*.*G610C*^ femora compared to control femora (Figure 5C) at 6 weeks of age but a significant reduction at 9 weeks (Figure 5G). This suggests that the genotype-dependent reduction in bone strength at 6 weeks can be explained by the smaller size but the bone strength defect persists in 9 week old mice due to the progressing material defect. This indicates that the OI phenotype progressively worsens between 6 and 9 weeks of age.

CBZ had no significant effect on any parameter measured by three point bending tests in *Col1a2* ^*+/p*.*G610C*^ mice at 6 or 9 weeks of age (Figure 5, Table S1, S2). However, in control mice, although 3 weeks of CBZ treatment did not modify bone strength, after 6 weeks of treatment maximum load, yield load and displacement were significantly lower than in vehicle-treated controls (Figure 5A, E and Table S1, S2). When corrected for bone width, no strength parameters were significantly altered by either 3 or 6 weeks of CBZ treatment (Figure 5C, D, G, H and Tables S1, S2). This indicates that CBZ treatment of 3 week old control mice for 6 weeks suppresses bone growth sufficiently to reduce functional strength.

## Discussion

This study was based on the premise that stimulation of autophagic mutant collagen I degradation by carbamazepine (CBZ) treatment might provide therapeutic benefit in OI. In collagen misfolding disorders, the best evidence for this approach comes from a mouse model of Metaphyseal Chondrodysplasia, Type Schmid (MCDS), where CBZ treatment stimulated autophagic degradation of the mutant misfolded collagen X, reducing ER stress and improving the cartilage and bone disease severity ^11,12^. This provided a strong rationale for testing the possible therapeutic effect of carbamazepine in the *Col1a2* ^*+/p*.*G610C*^ mouse model of OI. However, we found that treatment of 3 week-old *Col1a2* ^*+/p*.*G610C*^ mice for either 3 or 6 weeks with CBZ, at the dose effective in the MCDS model, had no therapeutic benefit for the deficiencies in bone structure, strength, or composition in *Col1a2* ^*+/p*.*G610C*^ mice. The study also provides new information about the *Col1a2* ^*+/p*.*G610C*^ mouse bone phenotype and reveals that CBZ treatment in mice with healthy skeletons can be detrimental.

The lack of beneficial effect of CBZ on *Col1a2* ^*+/p*.*G610C*^ bone is not due to unavailability of CBZ in the OI bone with this treatment regime, since the impact of CBZ on the control bone (discussed below) clearly indicates that the CBZ was bio-available in these growing bones. Furthermore, this dose of CBZ was effective in modifying the ER stress response of the MCDS mouse growth plate cartilage ^11,12^. This suggests that CBZ does not stimulate sufficient intracellular clearance of mutant misfolded/aggregated collagen I *in vivo* to have a positive clinical effect on downstream pathological signalling in the *Col1a2* ^*+/p*.*G610C*^ bone. Furthermore, while both MCDS and OI (*Col1a2* ^*+/p*.*G610C*^) are caused by mutant collagen misfolding and intracellular retention, the collagen types (collagen X and collagen I, respectively), the nature and context of the mutations are different ^6,10^, perhaps requiring different strategies to remove accumulated protein from the affected cells.

This is the second study showing that autophagy-stimulating drugs do not improve bone structure or strength in the *Col1a2* ^*+/p*.*G610C*^ mouse. Other approaches, based on clearing mutant collagen may still be effective, since rapamycin ^9^ and CBZ have known “off-target’ effects. An alternative approach that has gained traction in several diseases, is to reduce the mutant misfolded protein load, not by degradation, but by chemical chaperones such as 4-phenylbutyric aid (4-PBA) to assist in refolding the mutant proteins ^21^. 4-PBA reduces ER stress *in vitro* in fibroblasts with a range of mutations, including *COL1A1* and *COL1A2* misfolding mutations ^22,23^. *In vivo*, 4-PBA treatment improved bone abnormalities in a zebrafish collagen misfolding OI model ^24^, and in the *Aga2*^+/-^ mouse model, where a *Col1a1* mutations results in failed procollagen I assembly ^25^. While 4-PBA may not be the ideal drug because of multiple other effects, such as acting as an ammonia scavenger, histone deacetylase inhibitor and with effects on mitochondria and peroxisomes, it does show proof-of-principle that facilitating mutant collagen refolding could be a useful approach to further explore with different OI collagen I mutations.

The disappointing lack of clinical efficacy of CBZ in OI is, however, moot since we found that the 6 week CBZ treatment of 3 week-old growing bone in control mice was deleterious. CBZ impaired bone growth, in both longitudinal and transverse directions, and reduced the cortical and trabecular bone mass of healthy mice leading to a significant reduction in functional bone strength. Although CBZ treatment changed the morphology of the control bone to reach similar proportions to the OI bone, the strength defect caused by CBZ was less severe than the OI phenotype. This was because CBZ changed bone morphology without accompanying changes in bone composition, measured by sFTIRM.

CBZ is a widely used anti-epileptic drug (AED) and there have been multiple reports that short-and long-term CBZ treatment in children and adults with epilepsy leads to low serum 25-hydroxyvitamin D, and high serum parathyroid hormone and alkaline phosphatase ^26–28^. This suggests CBZ, like other anti-convulsant therapies, causes osteomalacia and secondary hyperparathyroidism and may increase fracture risk ^29^. However, analyses of the effect of CBZ on bone structure in children with epilepsy have been limited to small observational studies, with some reporting reduced areal bone mineral density (BMD) ^27,30^ but others indicating no significant effect ^31–34^. No studies have reported how CBZ in children influences bone growth. Pre-clinical studies have been limited to adult rats, and these too have conflicting results, with one study reporting no change in cortical or trabecular structure by micro-CT, nor any change in bone strength ^35^, but another reporting a significant decline in both trabecular bone mass and BMD at multiple sites ^36^. It will be important to systematically evaluate any effects of CBZ on normal bone structure and biomechanics to resolve the conflicting data and inform on the suitability of CBZ as a potential OI therapy.

This study also provides new information about the OI phenotype. Firstly, it shows that this phenotype can be detected as early as 6 weeks of age, a time point not previously studied, and that the defect in bone strength becomes more severe by 9 weeks of age. Our data suggests that the phenotype progression between 6 and 9 weeks of age is largely due to a worsening material defect. There are three pieces of evidence for this: (1) the difference in cortical bone size and shape between control and *Col1a2* ^*+/p*.*G610C*^ bones did not worsen with age, (2) the defective mineral:matrix ratio did become more severe with age, and (3) the worsening difference in bone strength persists after correcting for bone size.

Our work also provides new information about defects in skeletal maturation in the *Col1a2* ^*+/p*.*G610C*^ mouse. Bone undergoes multiple changes in post-embryonic growth. Not only do the bones lengthen, but cortical bone thickens, and the material changes from woven bone to an increasingly dense lamellar structure ^37,38^; this is reflected in the micro-CT scans showing a transition with increasing distance from the growth plate from a high proportion of low-density bone to a material containing more high-density bone ^13,16^. *Col1a2* ^*+/p*.*G610C*^ femora did exhibit cortical bone maturation, but the presence of less high density bone in the least mature region (close to the growth plate), and more low density bone in the most mature region (near the diaphysis), could indicate an initial more rapid process of cortical maturation and an early plateau, such that there is a higher proportion of mid-density bone along the full metaphyseal length. Our sFTIRM analysis also indicates that the greater proportion of mineral relative to collagen in *Col1a2* ^*+/p*.*G610C*^ bone is already significant at 6 weeks of age and is only detected in more mature regions of bone. This indicates that although mineral is initially deposited at normal levels on the periosteum of *Col1a2* ^*+/p*.*G610C*^ bone matrix, it accrues more rapidly in these mice than controls, probably driven by the defective collagen structure.

Our data raise two important considerations for any future applications of CBZ in genetic skeletal disease therapy. Firstly, given the lack of effect with this *Col1a2* ^*+/p*.*G610C*^ mutant, it seems unlikely that CBZ will be useful for broad-based OI therapy. It remains possible that a beneficial CBZ effect could be OI mutant-specific, but this would require extensive mutation-specific testing in patients. Secondly, and more importantly, the negative impact of CBZ on normal bone growth and structure is worrying in terms of any proposed therapies. If experimental evidence emerges that CBZ is beneficial with specific protein misfolding mutations, it will be critically important to closely monitor bone growth in clinical trials to ensure that any potential benefits outweigh the possible side-effects. This may be particularly important when treatment coincides with bone growth.

## Acknowledgments

This study was funded by an Australian National Health & Medical Research Council project grant (GNT 1049863) and Senior Research Fellowship (GNT 1154819) (to N.A.S.), philanthropic support to St. Vincent’s Institute, and the Victorian Government’s Operational Infrastructure Support Program. The FTIRM data was generated on the IRM beamline at the Australian Synchrotron, part of ANSTO, and we thank Pimm Vongsvivut and Mark Tobin for their assistance there; we also thank Mark Forwood, Griffith University, for consulting on the mechanical testing method.

## Statement of Author Contributions

JFB, SRL, NAS conceived the project. MB, JFB, SRL, NAS designed the experiments. MB, JFB, NAS analysed the data and wrote the manuscript, which was reviewed by all authors. MB, NEM, LR, LHWK, BC-I, ECW, JHG performed the experiments and analysed data.

## Conflict of Interest Statement

The authors declare no conflicts of interest

## Notes

### Competing Interest Statement

The authors have declared no competing interest.

