## Supplemental Data for "The effect of carbamazepine on bone structure and strength in control and Osteogenesis Imperfecta (*Col1a2* ^+/*p.G610C*^) mice"

**Figure S1. Multi-level-thresholding analysis along the femoral metaphysis showing low-, mid- and high-density bone tissue levels from 9 week old male control and *Col1a2* $^{+/p.G610C}$ mice treated with vehicle or carbamazepine (CBZ) for six weeks by micro-computed tomography (µCT).** Graphs shown describe genotype-dependent (A) and treatment-dependent (B, C) differences in bone densities. Data from control and *Col1a2* $^{+/p.G610C}$ mice from panel A are shown in panel B in panel C, respectively, with the effect of CBZ. Data are mean ± SEM; n= 5-8 mice/group. The line on top of the graph indicates a significant difference between the groups ($p<0.05$ was considered minimum significance).

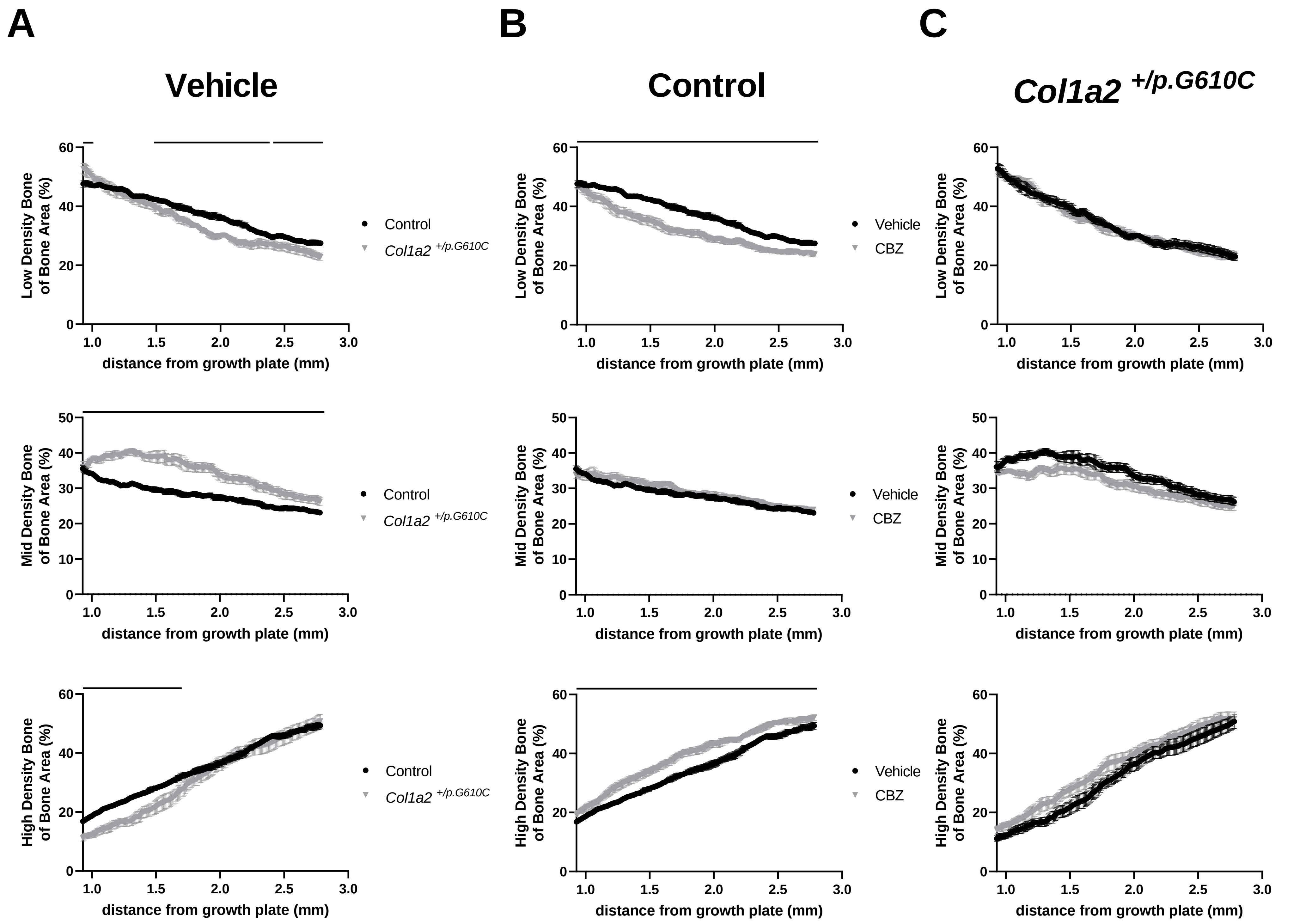
A
B
C
Vehicle
Control
Col1a2 +/p.G610C
Low Density Bone of Bone Area (%)
Mid Density Bone of Bone Area (%)
High Density Bone of Bone Area (%)
distance from growth plate (mm)
Control
Col1a2 +/p.G610C
Vehicle
CBZ

**Figure S2. Regional analysis of bone composition of tibial cortex from 6 week (A-D) and 9 week (E-H) old male control and *Col1a2* $^{+/p.G610C}$ mice treated with vehicle or carbamazepine (CBZ) for three and six weeks respectively assessed by synchrotron Fourier-transform infrared microspectroscopy (sFTIRM).** Ratios were calculated from integrated areas of phosphate (1180-916cm$^{-1}$), carbonate (890-852cm$^{-1}$), amide I (1712-1588cm$^{-1}$) and amide II (1600-1500cm$^{-1}$) curves. Crystallinity sub-peak was calculated by the integrated area from 1030-1020cm$^{-1}$. Data shown are mean ± SEM; n= 7-11 mice/group for 6 week old mice and n= 5-8 mice/group for 9 week old mice. + q<0.05, ++ q<0.01 +++ q<0.001 vs. genotype- and treatment-matched region 0 µm. # q<0.05, ## q<0.01, ### q<0.001 vs. treatment- and region-matched controls.

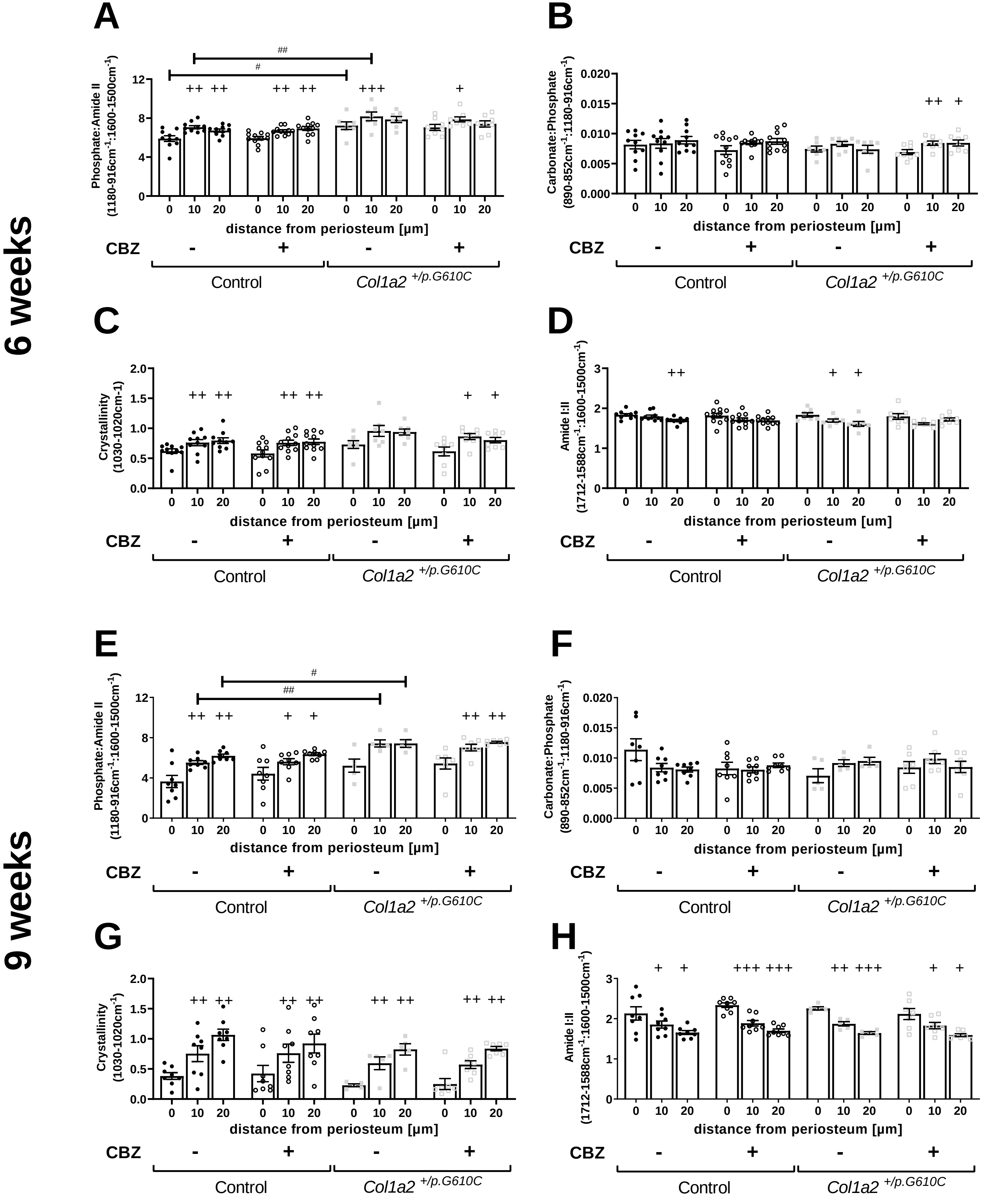

6 weeks
9 weeks
A
Phosphate:Amide II (1180-916cm-1:1600-1500cm-1)
##
#
++ ++ ++ ++ +++ +
B
Carbonate:Phosphate (890-852cm-1:1180-916cm-1)
++ +
C
Crystallinity (1030-1020cm-1)
++ ++ ++ ++ + +
D
Amide I:II (1712-1588cm-1:1600-1500cm-1)
++ + +
E
Phosphate:Amide II (1180-916cm-1:1600-1500cm-1)
#
##
++ ++ + + ++ ++
F
Carbonate:Phosphate (890-852cm-1:1180-916cm-1)
G
Crystallinity (1030-1020cm-1)
++ ++ ++ ++ ++ ++ ++ ++
H
Amide I:II (1712-1588cm-1:1600-1500cm-1)
+ + +++ +++ ++ +++ + +
0 10 20 0 10 20 0 10 20 0 10 20
distance from periosteum [µm]
CBZ - + - +
Control
Col1a2 +/p.G610C

**Table S1. Additional cortical bone dimensions assessed by µCT and bone strength parameters assessed by three point bending test of femora from 6 week old male control and *Col1a2* $^{+/p.G610C}$ mice treated with vehicle or carbamazepine (CBZ) for three weeks.** Data shown are mean ± SEM, n= 7-16 mice/group. * p<0.05, ** p<0.01, *** p<0.001 vs. treatment-matched controls.

### Bone dimensions

| | Control | | *Col1a2* $^{+/p.G610C}$ | |
|---|---|---|---|---|
| | Vehicle (n= 10) | CBZ (n= 11) | Vehicle (n= 7) | CBZ (n= 8) |
| Marrow Area ($mm^2$) | 1.07 ± 0.03 | 1.07 ± 0.02 | 0.88 ± 0.02 *** | 0.91 ± 0.02 |
| Mean Polar Moment of Inertia ($mm^4$) | 0.34 ± 0.03 | 0.28 ± 0.01 | 0.22 ± 0.02 *** | 0.21 ± 0.01 |
| Medio-lateral width (mm) | 1.72 ± 0.03 | 1.66 ± 0.02 | 1.53 ± 0.03 *** | 1.55 ± 0.02 |
| Cranio-caudal width (mm) | 1.30 ± 0.02 | 1.28 ± 0.01 | 1.23 ± 0.03 * | 1.20 ± 0.02 |

### Bone strength parameters

| | Control | | *Col1a2* $^{+/p.G610C}$ | |
|---|---|---|---|---|
| | Vehicle (n= 11) | CBZ (n= 16) | Vehicle (n= 7) | CBZ (n= 7) |
| **Structural properties** | | | | |
| Yield Load (N) | 8.09± 0.59 | 6.75 ± 0.26 | 6.01 ± 0.72 * | 5.29 ± 0.47 |
| Yield Displacement (mm) | 0.20 ± 0.02 | 0.18 ± 0.02 | 0.11 ± 0.01 * | 0.14 ± 0.02 |
| Stiffness (N/mm) | 45.9 ± 3.67 | 44.0 ± 4.60 | 51.2 ± 5.24 | 34.9 ± 2.92 |
| **Material properties** | | | | |
| Yield Stress (MPa) | 18.5 ± 1.12 | 19.6 ± 1.03 | 21.9 ±1.02 | 20.2 ± 0.95 |
| Yield Strain (%) | 4.29 ± 0.43 | 3.77 ± 0.38 | 2.33 ± 0.28 ** | 2.757 ± 0.33 |
| Elastic Modulus (MPa) | 467 ± 49.3 | 601 ± 77.9 | 936 ± 116 ** | 700 ± 86.2 |

**Table S2. Additional cortical bone dimensions assessed by µCT and bone strength parameters assessed by three point bending test of femora from 9 week old male control and *Col1a2* $^{+/p.G610C}$ mice treated with vehicle or carbamazepine (CBZ) for six weeks.** Data shown are mean ± SEM, n= 5-8 mice/group. * p<0.05, ** p<0.01, *** p<0.001 vs. treatment-matched controls. # p<0.05, ## p<0.01, ### p<0.001 vs. genotype-matched controls.

### Bone dimensions

| | **Control** | | ***Col1a2*** $^{+/p.G610C}$ | |
|---|---|---|---|---|
| | Vehicle (n= 10) | CBZ (n= 11) | Vehicle (n= 7) | CBZ (n= 8) |
| Marrow Area ($mm^2$) | 1.07 ± 0.03 | 1.07 ± 0.02 | 0.88 ± 0.02 *** | 0.91 ± 0.02 |
| Mean Polar Moment of Inertia ($mm^4$) | 0.34 ± 0.03 | 0.28 ± 0.01 | 0.22 ± 0.02 *** | 0.21 ± 0.01 |
| Medio-lateral width (mm) | 1.72 ± 0.03 | 1.66 ± 0.02 | 1.53 ± 0.03 *** | 1.55 ± 0.02 |
| Cranio-caudal width (mm) | 1.30 ± 0.02 | 1.28 ± 0.01 | 1.23 ± 0.03 * | 1.20 ± 0.02 |

### Bone strength parameters

| | **Control** | | ***Col1a2*** $^{+/p.G610C}$ | |
|---|---|---|---|---|
| | Vehicle (n= 11) | CBZ (n= 16) | Vehicle (n= 7) | CBZ (n= 7) |
| **Structural properties** | | | | |
| Yield Load (N) | 8.09± 0.59 | 6.75 ± 0.26 | 6.01 ± 0.72 * | 5.29 ± 0.47 |
| Yield Displacement (mm) | 0.20 ± 0.02 | 0.18 ± 0.02 | 0.11 ± 0.01 * | 0.14 ± 0.02 |
| Stiffness (N/mm) | 45.9 ± 3.67 | 44.0 ± 4.60 | 51.2 ± 5.24 | 34.9 ± 2.92 |
| **Material properties** | | | | |
| Yield Stress (MPa) | 18.5 ± 1.12 | 19.6 ± 1.03 | 21.9 ±1.02 | 20.2 ± 0.95 |
| Yield Strain (%) | 4.29 ± 0.43 | 3.77 ± 0.38 | 2.33 ± 0.28 ** | 2.757 ± 0.33 |
| Elastic Modulus (MPa) | 467 ± 49.3 | 601 ± 77.9 | 936 ± 116 ** | 700 ± 86.2 |
